# Systematic analysis of naturally occurring insertions and deletions that alter transcription factor spacing identifies tolerant and sensitive transcription factor pairs

**DOI:** 10.1101/2020.04.02.021535

**Authors:** Zeyang Shen, Rick Z. Li, Thomas A. Prohaska, Marten A. Hoeksema, Nathan J. Spann, Jenhan Tao, Gregory J. Fonseca, Thomas Le, Lindsey Stolze, Mashito Sakai, Casey E. Romanoski, Christopher K. Glass

## Abstract

Regulation of gene expression requires the combinatorial binding of sequence-specific transcription factors (TFs) at promoters and enhancers. Prior studies showed that alterations in the spacing between TF binding sites can influence promoter and enhancer activity. However, the relative importance of TF spacing alterations resulting from naturally occurring insertions and deletions (InDels) has not been systematically analyzed. To address this question, we first characterized the genome-wide spacing relationships of 75 TFs in K562 cells as determined by ChIP-sequencing. We found a dominant pattern of a relaxed range of spacing between collaborative factors, including 46 TFs exclusively exhibiting relaxed spacing with their binding partners. Next, we exploited millions of InDels provided by genetically diverse mouse strains and human individuals to investigate the effects of altered spacing on TF binding and local histone acetylation. Spacing alterations resulting from naturally occurring InDels are generally tolerated in comparison to genetic variants directly affecting TF binding sites. A remarkable range of tolerance was further established for PU.1 and C/EBPβ, which exhibit relaxed spacing, by introducing synthetic spacing alterations ranging from 5-bp increase to >30-bp decrease using CRISPR/Cas9 mutagenesis. These findings provide implications for understanding mechanisms underlying enhancer selection and for the interpretation of non-coding genetic variation.

## Introduction

Genome-wide association studies (GWASs) have identified thousands of genetic variants associated with diseases and other traits (MacArthur et al., 2017; Visscher et al., 2017). Single nucleotide polymorphisms (SNPs) and short insertions and deletions (InDels) represent common forms of these variants. The majority of GWAS variants fall at non-protein-coding regions of the genome, implicating their effects on gene regulation (Farh et al., 2015; Ward & Kellis, 2012). Gene expression is regulated by transcription factors (TFs) in a cell-type-specific manner. TFs bind to short, degenerate sequences at promoters and enhancers, often referred to as TF binding motifs. Active promoters and enhancers are selected by combinations of sequence-specific TFs that bind in an inter-dependent manner to closely spaced motifs. SNPs and InDels can create or disrupt TF binding motifs and are a well-established mechanism for altering gene expression and biological function (Behera et al., 2018; Deplancke et al., 2016; Grossman et al., 2017; Heinz et al., 2013). InDels can additionally change spacing between motifs, but it remains unknown the extent to which altered spacing is relevant for interpreting genetic variation in human populations or between animal species.

Previous studies reported two major categories of motif spacing between inter-dependent TFs (Slattery et al., 2014). One category refers to the enhanceosome model (Slattery et al., 2014) that requires specific or “constrained” spacing. It is mainly provided by TFs that form ternary complexes recognizing composite binding motifs, exemplified by GATA, Ets and E-box transcription factors in mouse hematopoietic cells (Ng et al., 2014), MyoD and other cell-type-specific factors in muscle cells (Nandi et al., 2013), Sox2 and Oct4 in embryonic stem cells (Rodda et al., 2005). In vitro studies of the binding of pair-wise combinations of ~100 TFs to a diverse library of DNA sequences identified 315 out of 9400 possible interactive TF pairs that select composite elements with constrained positions of the respective recognition motifs (Jolma et al., 2015). Constrained spacing required for the optimal binding and function of interacting TFs can also occur between independent motifs, such as occurs at the interferon-β enhanceosome (Panne, 2008). In comparison to constrained spacing, another category of motif spacing allows TFs to interact over a relatively broad range (e.g., 100-200 bp), which we call “relaxed” spacing and is equivalent to the billboard model (Slattery et al., 2014). This type of spacing relationship is observed in collaborative or co-occupied TFs that do not target promoters or enhancers as a ternary complex (Heinz et al., 2010; Jiang & Singh, 2014; Sönmezer et al., 2021).

Substantial evidence indicates that the two categories of spacing requirement can experience a different level of impact from genetic variation. Reporter assays examining synthetic alterations of motif spacing revealed examples of TFs that require constrained spacing and have high sensitivity of transcription factor binding and gene expression on spacing (Farley et al., 2015; Ng et al., 2014; Panne, 2008). On the contrary, flexibility in motif spacing has been demonstrated using reporter assays in Drosophila (Menoret et al., 2013) and HepG2 cells (Smith et al., 2013).

However, these studies did not distinguish the impact of altered spacing on transcription factor binding or subsequent recruitment of co-activators required for gene activation. Moreover, it remains unknown the extent to which these findings are relevant to spacing alterations resulting from naturally occurring genetic variation.

To investigate the effects of altered spacing on TF binding and function, we first characterized the genome-wide binding patterns of seventy-five TFs based on their binding sites determined by chromatin immuno-precipitation sequencing (ChIP-seq). We developed a computational framework that assigned each spacing relationship to “constrained” or “relaxed” category and associated spacings to the naturally occurring InDels observed in human populations to study the selective constraints of different spacing relationships. As specific case studies, we leveraged natural genetic variation in numerous human samples and from five strains of mice to study the effect size of spacing alterations on TF binding activity and local histone acetylation. We find that InDels altering spacing are generally less constrained and well tolerated when they occur between TF pairs with relaxed spacing relationships. Finally, we established remarkable tolerance in spacing for macrophage lineage determining TFs (LDTFs), PU.1 and C/EBPβ, by introducing a wide range of InDels between their respective binding sites at representative endogenous genomic loci using CRISPR/Cas9 mutagenesis in mouse macrophages.

## Results

### Transcription factors primarily co-bind with relaxed spacing

We characterized spacing relationships for 75 TFs of K562 cells covering diverse TF families (Hu et al., 2019) based on the ChIP-seq data from ENCODE data portal (Davis et al., 2018). After obtaining reproducible TF binding sites, we first used the corresponding position weight matrix (PWM) of each TF (Figure 1-table supplement 1) to scan through the sequence of every binding site and identified the locations of high-affinity motifs that are less than 50 bp from ChIP-seq peak centers (Fig. 1A; Figure 1-figure supplement 1). The binding sites of every pair of TFs were then merged to compute the edge-to-edge motif spacing at all the co-binding sites. Motif spacings were eventually aggregated to show a distribution within +/− 100 bp. To categorize spacing relationships, we used permutation tests on the gradients to test for specific spacing constraints and used Kolmogorov–Smirnov test (KS test) to test for a relaxed spacing relationship against random distribution.

**Figure 1.**
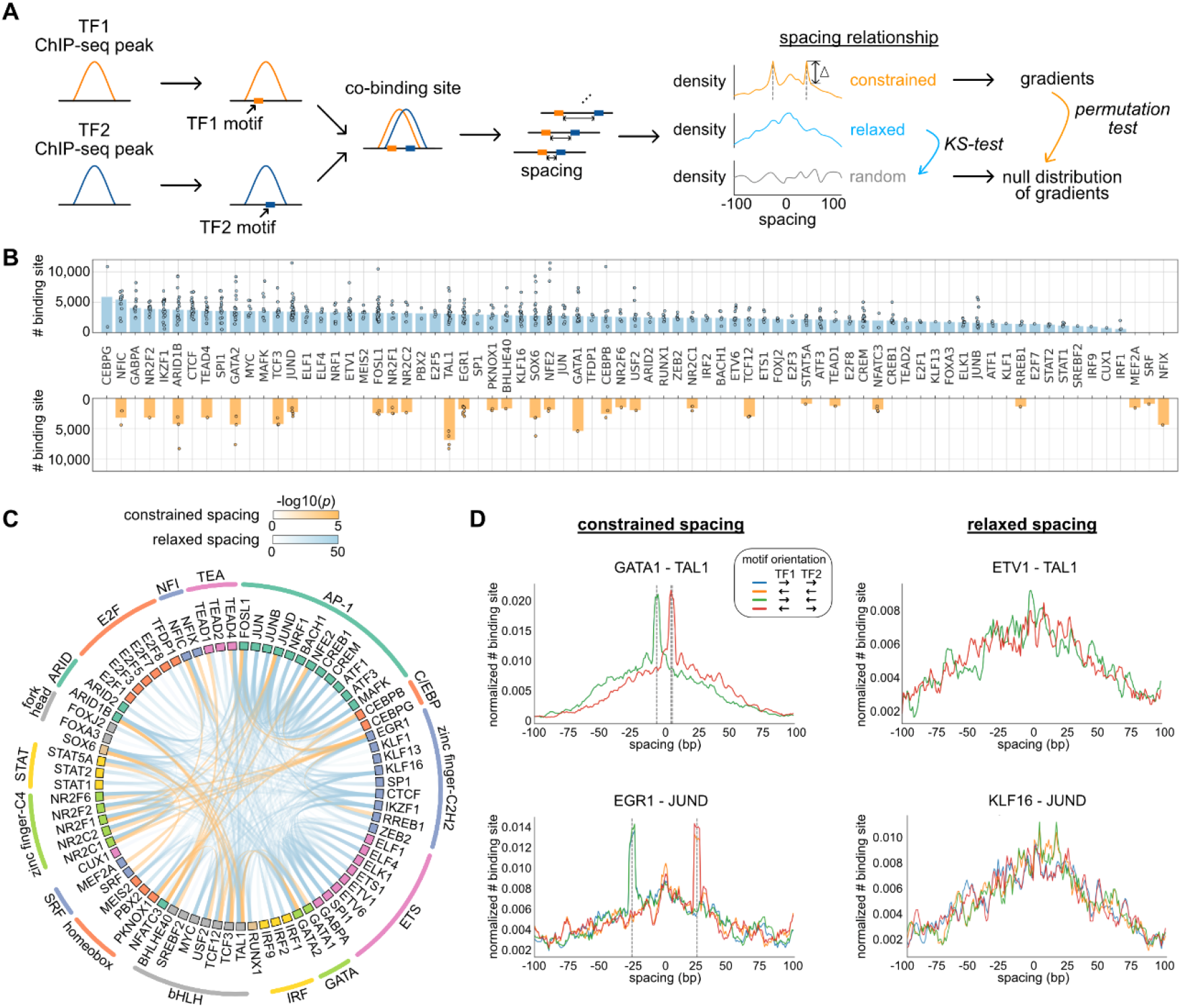
Characterization of spacing relationships for transcription factor pairs. (**A**) Schematic of data analysis pipeline for characterizing the spacing relationships based on TF ChIP-seq data. (**B**) Dissection of TF binding sites for TFs in K562 cells based on spacing relationships with co-binding TFs. Each dot represents the co-binding peak number of the corresponding TF and one of its co-binding TFs. Bar heights indicate means among the co-binding TFs. (**C**) Circos plot summarizing spacing relationships for all the TF pairs analyzed. Orange and blue bands represent significant constrained and relaxed spacing relationships, respectively. Color opacity indicates the level of significance. TFs are grouped and colored by TF family. (**D**) The spacing distributions of example TF pairs with constrained spacing or relaxed spacing relationships. Dashed lines indicate the significant constrained spacings. Since TAL1 motif is completely palindromic, the motif orientation is only differentiated by its co-binding partners.

We applied this computational framework to all possible pairs of TFs. By dissecting each TF’s binding sites based on their spacing relationships with co-binding TFs, we found that 46 of the 75 TFs examined exclusively exhibited relaxed spacing relationships with other TFs (Fig. 1B). 26 factors could participate in either relaxed or constrained interactions, depending on the specific co-binding TFs. Only 3 TFs interacted with only constrained spacing, some of which might show additional relaxed spacing relationships by expanding the current set of TFs. The significant pairwise patterns of relaxed and constrained spacing relationships are illustrated in Figure 1C. Among 32 TF pairs with constrained spacing relationships, most bind closely to each other within 15 bp spacing (Figure 1-figure supplement 2; Figure 1-table supplement 2). Some of these TF pairs have been reported to recognize composite motifs such as GATA1-TAL1 and NFATC3-FOSL1 (Macián et al., 2001; Ng et al., 2014) (Fig. 1D; Figure 1-figure supplement 3), and some are novel constrained spacing patterns discovered by our analysis such as MEF2A-JUND and CEBPB-TEAD4 (Figure 1-figure supplement 3). There are also TF pairs, exemplified by EGR1 and JUND, that bind relatively further away from each other but still require constrained spacing (Fig. 1D). Previous studies demonstrated interactions between EGR1 and AP-1 factors (Levkovitz & Baraban, 2002; Nakashima et al., 2003), but the underlying mechanism for such constrained spacing at 29 bp needs to be further investigated. TFs exhibiting relaxed spacing are exemplified by ETV1-TAL1 and JUND-KLF16, in which the frequency of co-binding progressively declines with distance from the center of the reference TF (Fig. 1D). In addition, the same type of spacing relationship is usually observed in different motif orientations (Fig. 1D), consistent with previous findings (Lis & Walther, 2016). The same TF pairs can have similar spacing relationships in different cell types, exemplified by CEBPB and JUND in K562 and HepG2 cells (Figure 1-figure supplement 4).

### Natural genetic variants altering spacing between relaxed transcription factors are associated with less deleteriousness in human populations

Based on a global view of the TF spacing relationships, we then studied whether these relationships associate with different levels of sensitivity to spacing alterations. Here, we leveraged more than 60 million InDels from gnomAD data (Karczewski et al., 2020), which were based on more than 75,000 genomes from unrelated individuals. We overlaid these InDels at motifs, between motifs, or within background regions of representative TF pairs of constrained and relaxed spacing relationships. We found that InDels at different regions have relatively similar distributions of InDel sizes with the majority being less than 5 bp (Fig. 2A). Next, we divided these InDels based on the allele frequency (AF) and the allele count (AC) into high-frequency variants (AF > 0.01%), rare variants (AF < 0.01%, AC > 1), and singletons (AC = 1). Most of the InDels at TF binding sites are singletons or rare variants (Figure 2-figure supplement 1). We compared the enrichment of these categories of InDels between TFs with different spacing relationships (Fig. 2B). The InDel compositions at motifs were not significantly different between constrained and relaxed spacing groups. On the contrary, singletons were significantly more enriched between motifs with constrained spacing, whereas high-frequency variants were significantly more depleted between these motifs. We also computed for several TF pairs with random spacing relationships as negative controls and found similar enrichments of InDels like those with relaxed spacing. Since common variants are associated with less deleteriousness and rare variants with more deleteriousness (Lek et al., 2016), our data suggest that InDels between motifs of TFs with constrained spacing could be just as damaging as those at motifs whereas InDels between motifs of TFs with relaxed spacing might have a much weaker effect.

**Figure 2.**
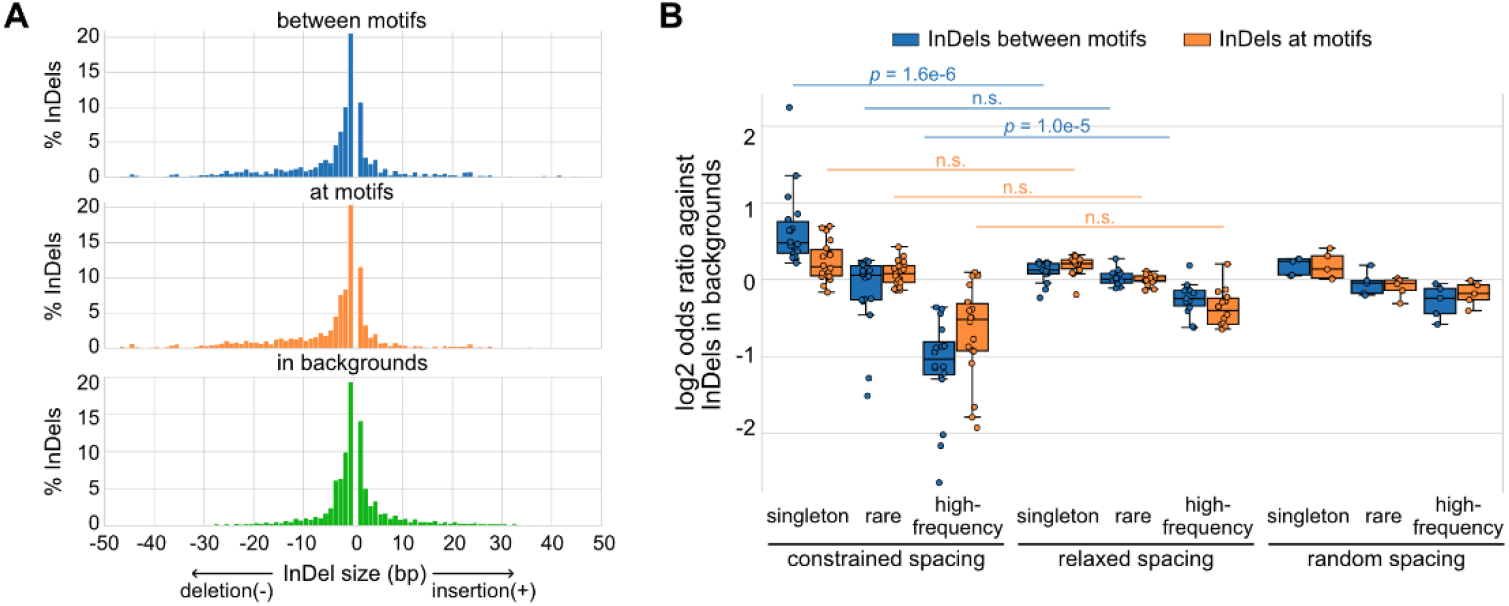
Naturally occurring InDels in human populations. (**A**) Size distributions of human InDels within different regions. (**B**) Log2 odds ratios for different categories of InDels. Each dot represents a TF pair with corresponding spacing relationship. Mann-Whitney U test was used to compare the odds ratios between different spacing relationships.

### Spacing alterations across mouse strains are generally tolerated by relaxed transcription factor binding and promoter and enhancer function

To investigate the regulatory effects of naturally occurring InDels that alter motif spacing, we leveraged more than 50 million SNPs and 5 million InDels from five genetically diverse mouse strains and the ChIP-seq data of key TFs and histone acetylation for the bone marrow-derived macrophages (BMDMs) from every mouse strain (Link, Duttke, et al., 2018). The five mouse strains include C57BL/6J (C57), BALB/cJ (BALB), NOD/ShiLtJ (NOD), PWK/PhJ (PWK), and SPRET/EiJ (SPRET).We first characterized the spacing relationship between the macrophage lineage-determining TFs (LDTFs) PU.1 and C/EBPβ, which have been found to bind in a collaborative manner at regulatory regions of macrophage-specific genes (Heinz et al., 2010). Based on our computational framework for characterizing spacing relationships (Fig. 1A), these two TFs follow a relaxed spacing relationship independent of their motif orientations (Fig. 3A; KS p-value < 1e-6). Moreover, both PU.1 and C/EBPβ binding activities quantified by the ChIP-seq tags were not correlated with motif spacing, suggesting no direct association between spacing and TF binding (Fig. 3B).

**Figure 3.**
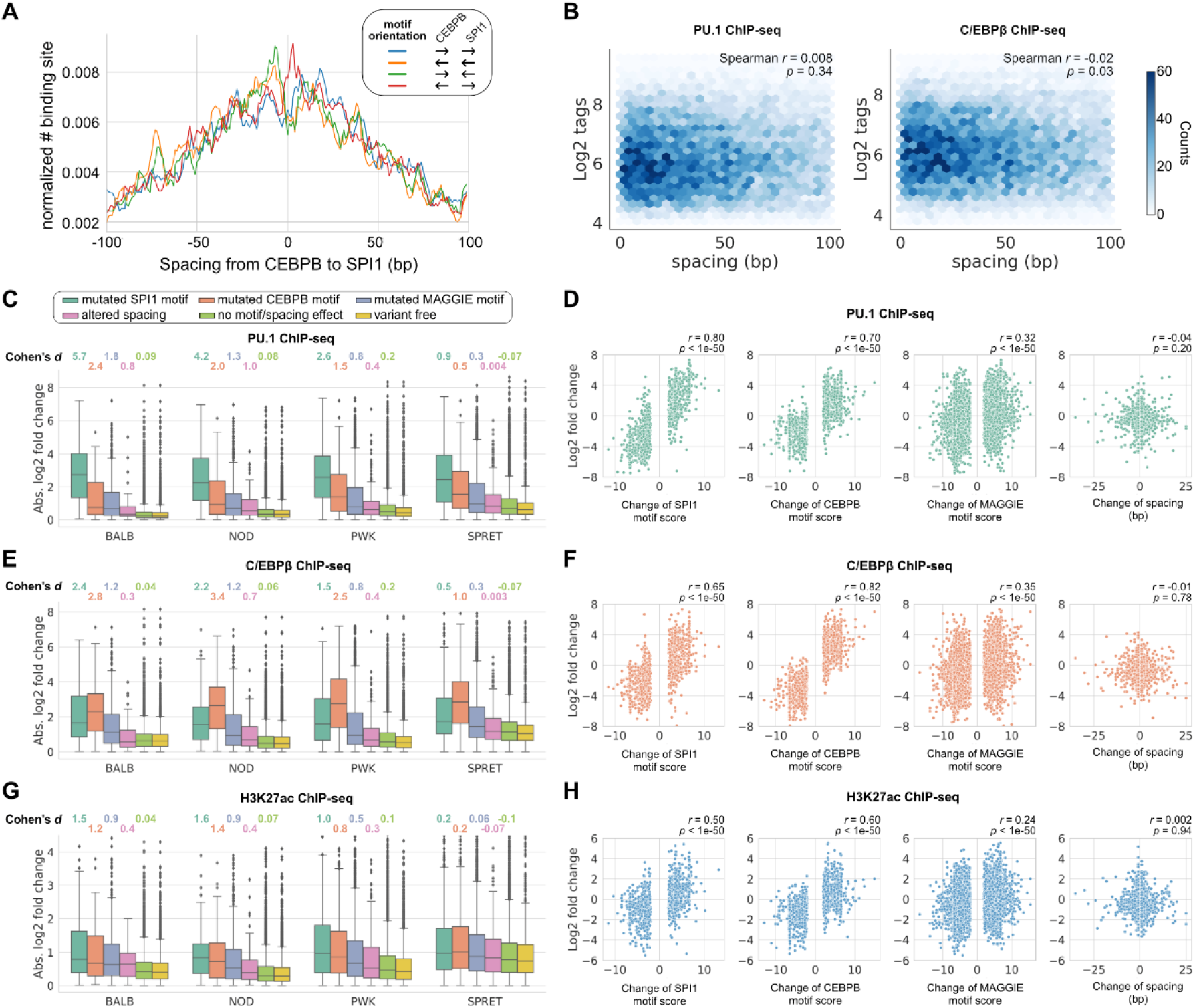
Effects of spacing alterations resulting from natural genetic variation across mouse strains. (**A**) Spacing distributions of PU.1 and C/EBPβ motif at co-binding sites. (**B**) Density plots showing the relationship between TF binding activity and motif spacing for the co-binding sites. Log2 ChIP-seq tags were calculated within 300 bp to quantify the binding activity of PU.1 and C/EBPβ. The color gradients represent the number of sites. Spearman correlation coefficients together with p-values are displayed. (**C, E, G**) Absolute log2 fold changes of ChIP-seq tags between C57 and another strain for (**C**) PU.1 binding, (**E**) C/EBPβ binding, or (**G**) H3K27ac level. Boxplots show the median and quartiles of every distribution. Cohen’s d effect sizes comparing against variant-free regions are displayed on top. (**D, F, H**) Correlations between change of motif spacing or motif score and change of (**D**) PU.1 binding, (**F**) C/EBPβ binding, or (**H**) H3K27ac level. Spearman correlation coefficients together with p-values are displayed.

We then conducted independent comparisons between C57 and one of the other four strains to investigate the effects of spacing alterations caused by natural genetic variation, which are mostly less than 5 bp like natural InDels in human populations (Figure 3-figure supplement 1). We first identified the co-binding sites of PU.1 and C/EBPβ for each strain and then, for each pairwise analysis, pooled the co-binding sites of C57 and the compared strain to obtain the testing set of regions. Based on the impacts of genetic variants on motif affinity and motif spacing, we categorized the testing regions into the following non-overlapping groups: 1) mutated PU.1 (i.e., SPI1) motif, 2) mutated C/EBPβ (i.e., CEBPB) motif, 3) mutated other functional motifs (i.e., MAGGIE motif), 4) altered spacing, 5) no motif affinity/spacing effect, and 6) variant free. Functional motifs were identified from PU.1 and C/EBPβ binding sites separately using MAGGIE (Shen et al., 2020), which is a computational tool that can identify motifs whose affinity changes are associated with TF binding changes (Figure 3-figure supplement 2). The effect of genetic variation was quantified by the log2 fold difference of ChIP-seq tag counts between strains at orthogonal sites (Fig. 3C). All the four independent comparisons showed that PU.1 binding is most strongly affected by PU.1 motif mutation, followed by C/EBPβ motif mutation and other functional motif mutation. Spacing alterations have a smaller effect size than any of these motif mutations, but still a relatively larger effect than variants affecting neither motif affinity nor spacing. Despite the moderate effect size of spacing alterations, we found such effect was independent of the size or direction of InDels (Fig. 3D). On the contrary, changes of PU.1 ChIP-seq tags are strongly correlated with changes of motif affinity (Fig. 3D). In addition, the effects of motif mutation and spacing alteration are not varied by the initial spacing between PU.1 and C/EBPβ motifs (Figure 3-figure supplement 3). Similar findings were observed in C/EBPβ binding, except that expectedly C/EBPβ motif mutation had the largest effect size and the strongest correlation with C/EBPβ binding activity (Fig. 3E, F; Figure 3-figure supplement 3).

To investigate whether the effects of altered spacing on PU.1 and C/EBPβ binding can be generalized to hierarchical interactions with signal-dependent transcription factors (SDTFs), we leveraged the ChIP-seq data of PU.1, the NFκB subunit p65, and an AP-1 factor cJun for BMDMs treated with the TLR4-specific ligand Kdo2 lipid A (KLA) in the same five strains of mice (Link, Duttke, et al., 2018). Upon macrophage activation with KLA, p65 enters the nucleus and primarily binds to poised enhancer elements that are selected by LDTFs including PU.1 and AP-1 factors (Heinz et al., 2015). We observed a relaxed spacing relationship between PU.1 and p65 and between cJun and p65 (Figure 3-figure supplement 4). In addition, InDels altering motif spacing had a much smaller effect size on TF binding than motif mutations (Figure 3-figure supplement 5), consistent with our findings from PU.1 and C/EBPβ.

Although alterations in motif spacing had generally weak effects at the level of DNA binding, it remained possible that changes in motif spacing could influence subsequent steps in enhancer and promoter activation. To examine this, we extended our analysis to local acetylation of histone H3 lysine 27 (H3K27ac), which is a histone modification that is highly correlated with enhancer and promoter function (Creyghton et al., 2010). We leveraged the H3K27ac ChIP-seq data of untreated BMDMs in the five strains of mice (Link, Duttke, et al., 2018) and calculated the log fold changes of H3K27ac level within the extended 1,000-bp regions of the PU.1 and C/EBPβ co-binding sites. Like for TF binding, altered spacing demonstrated weaker effects on histone acetylation than motif mutations (Fig. 3G; Figure 3-figure supplement 3), which is supported by the high consistency between change of TF binding and change of histone acetylation (Figure 3-figure supplement 6). The relative tolerance of spacing alteration was further reflected by a weak correlation between the change of acetylation level and the size of InDels, in comparison to a much stronger correlation with changes in motif affinity (Fig. 3H).

### Human quantitative trait loci altering spacing between relaxed transcription factors have small effect sizes

To study the effects of spacing alteration on transcription factor binding and local histone acetylation in human cells, we leveraged the ChIP-seq data of ERG, p65, and H3K27ac in endothelial cells from dozens of individuals (Stolze et al., 2020). ERG is an ETS factor that functions as an LDTF in endothelial cells that selects poised enhancers where p65 binds in a hierarchical manner upon interleukin-1β (IL-1β) stimulation (Hogan et al., 2017). ERG and p65 follow a relaxed spacing relationship according to our method (Fig. 4A). Next, we obtained 557 TF binding quantitative trait loci (bQTLs) for ERG, 5,791 bQTLs for p65, 25,621 histone modification QTLs (hQTLs) for H3K27ac in untreated cells, and 21,635 hQTLs for H3K27ac in IL-1β-treated cells (Stolze et al., 2020). We further classified bQTLs and hQTLs based on their impacts on motif affinity and spacing: 1) mutated both ERG and p65 (i.e., RELA) motif, 2) mutated ERG motif only, 3) mutated p65 motif only, 4) mutated other functional motifs identified by MAGGIE (Shen et al., 2020), 5) altered spacing between ERG and p65 motif, 6) none of the above. To find functional motifs, we fed MAGGIE with 100-bp sequences around QTLs before and after swapping alleles at the center (Figure 4-figure supplement 1). As a result, only a small portion of bQTLs and hQTLs directly mutates an ERG or RELA motif (Fig. 4B; Figure 4-figure supplement 2). However, such motif mutations are enriched in bQTLs compared to non-QTLs (Fisher’s exact p < 1e-4). On the contrary, InDels that alter motif spacing are significantly depleted in p65 bQTLs (Fisher’s exact p = 1.3e-15). These InDels from the dozens of individuals are predominantly shorter than 5 bp by following a similar size distribution of those in human populations (Figure 4-figure supplement 3). A large proportion of QTLs affect other functional motifs, implicating the complexity of TF interactions. More than a quarter of the QTLs affect neither motif affinity nor motif spacing, which can be explained by the high correlation of non-functional variants with functional variants due to linkage disequilibrium.

**Figure 4.**
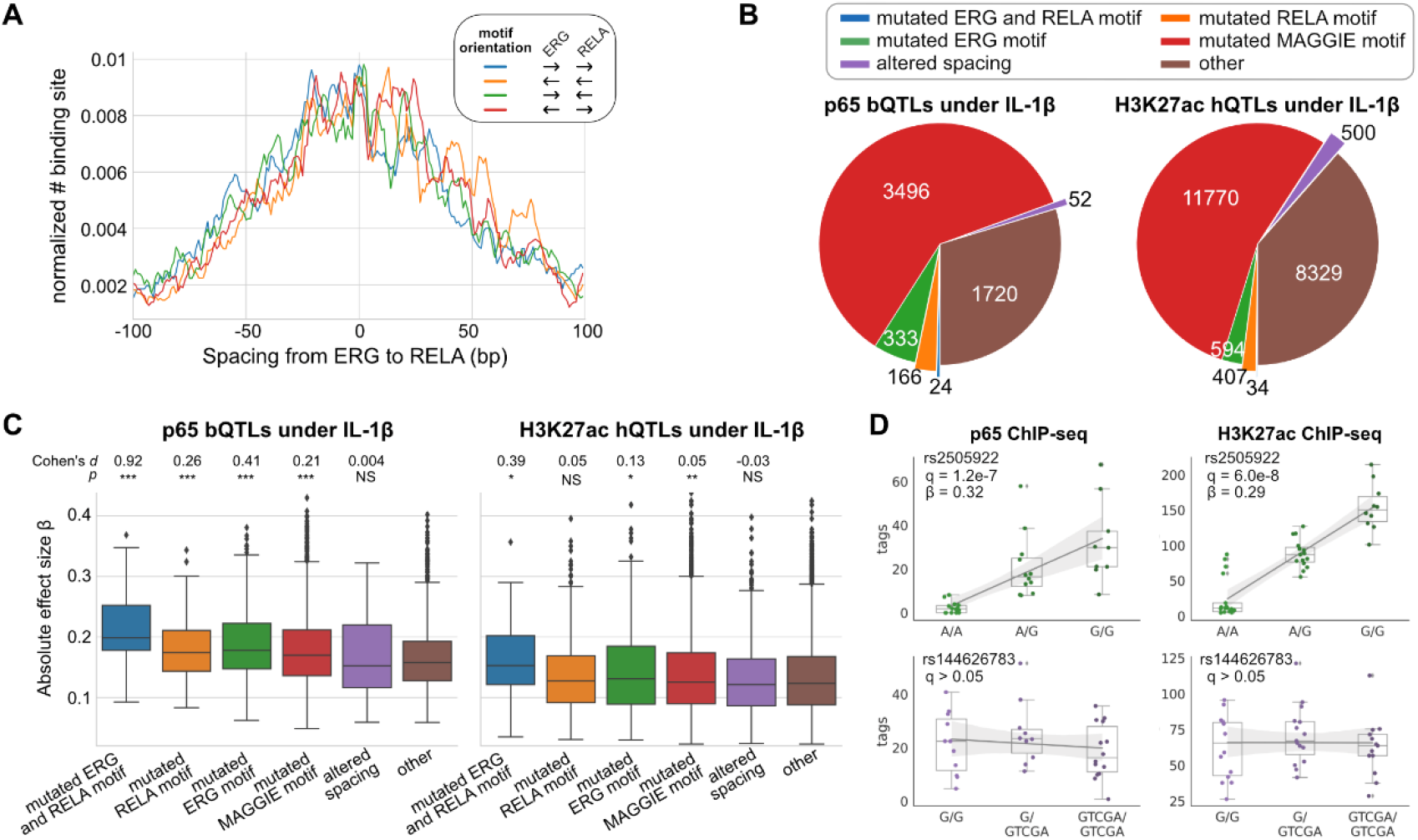
Effects of chromatin QTLs in human endothelial cells. (**A**) Spacing distributions of ERG and RELA motif at co-binding sites. (**B**) Classification of chromatin QTLs based on the impacts on motif and spacing. (**C**) Absolute correlation coefficients of different QTLs. Cohen’s d and Mann-Whitney U test p-values comparing against the “other” group are displayed on top. *p<0.01, **p<0.001, ***p<0.0001. (**D**) Example QTLs for large effect size due to ERG motif mutation (upper) and trivial effect due to spacing alteration (lower).

We further compared the effect sizes of different categories of QTLs. Despite being the minority among QTLs, variants that mutate both ERG and RELA motifs have the strongest effects on both p65 binding and histone acetylation in IL-1β-treated endothelial cells (Fig. 4C). In comparison, ERG binding and the basal level of histone acetylation are significantly affected by ERG motif mutations in untreated endothelial cells and not by p65 motif mutations, consistent with the hierarchical interaction of p65 only upon IL-1β stimulation (Figure 4-figure supplement 4). In both conditions of endothelial cells, spacing alterations have the smaller effect size than motif mutation categories and are not significantly different from likely non-functional variants in the “other” group. The examples showed a variant being both a p65 bQTL and a H3K27ac hQTL under the IL-1β state due to its impact on an ERG motif, and a 4-bp insertion between ERG and p65 motifs associated with no change in p65 binding or H3K27ac (Fig. 4D).

### Relaxed transcription factor binding is highly tolerant to synthetic spacing alterations

The generally small effects of InDels occurring between TF pairs exhibiting relaxed spacing relationships raised the question of the robustness and the extent of such tolerance at genomic locations lacking such variation. We addressed this question by using CRISPR/Cas9 editing to introduce synthetic InDels between binding sites observed for the LDTFs PU.1 and C/EBPβ in mouse macrophages (Fig. 5A). We used lentiviral transduction in Cas9-expressing ER-HoxB8 cells, which are conditionally immortalized monocyte progenitors, to introduce gRNAs targeting genomic sequences between the locations of PU.1 and C/EBPβ co-binding. The Cas9 nuclease activity at these sites resulted in non-homologous DNA repair that generated various sizes of InDels in the populations of transduced cells. After sorting the successfully transduced ER-HoxB8 cells and differentiating them into macrophages, we performed ChIP for C/EBPβ and deeply sequenced amplicons of the target regions of Cas9 cleavage. Lastly, the reads were mapped to the target regions by allowing various sizes of gaps at the cut sites and were quantified by comparing to the input DNA samples.

**Figure 5.**
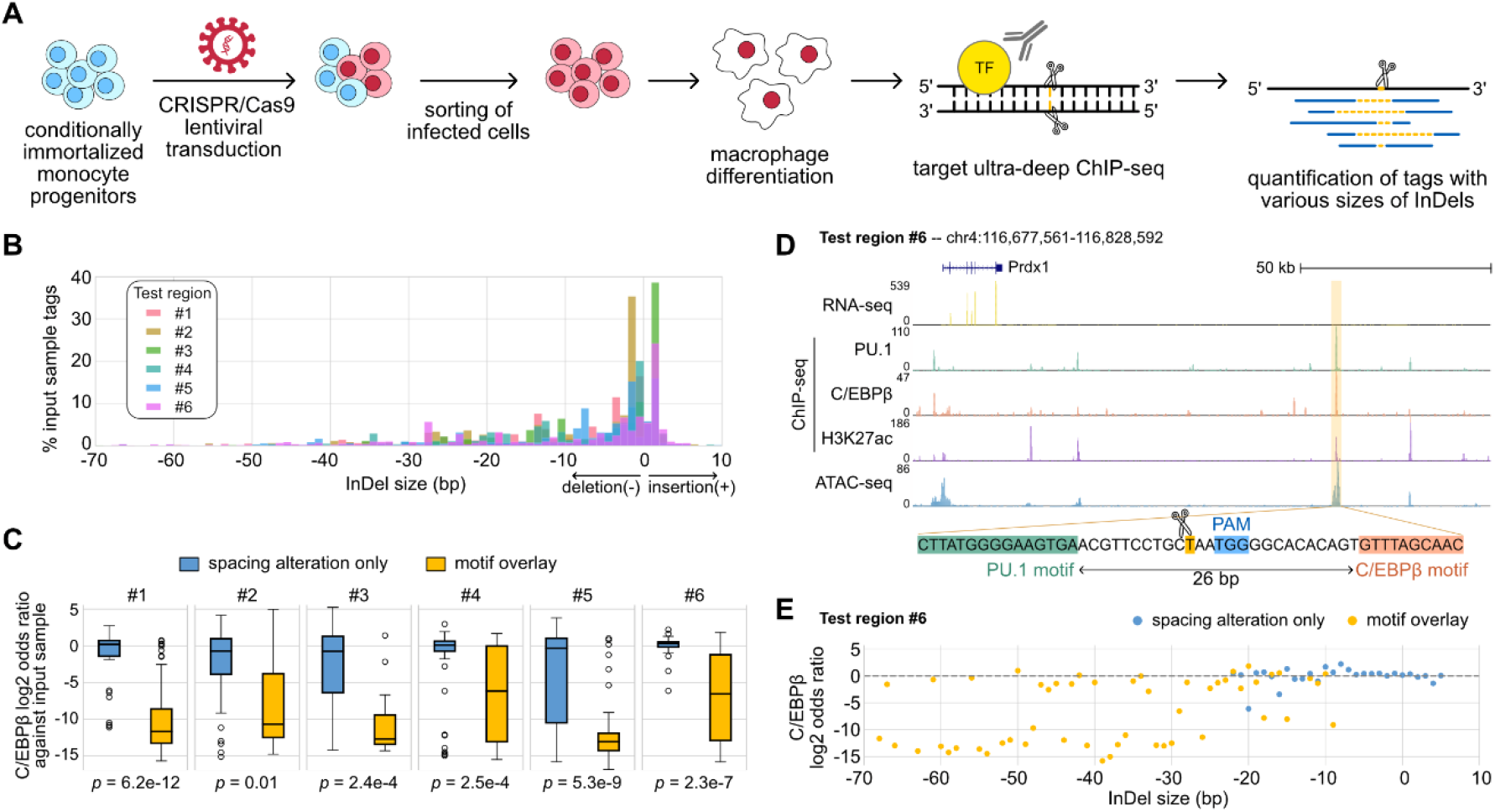
Effects of variable sizes of synthetic spacing alterations. (**A**) Schematic for generating and analyzing synthetic spacing alterations. (**B**) The distributions of valid read counts from the input sample based on the InDel sizes of the reads. Negative InDel size indicates deletion, and positive size means insertion. (**C**) Log2 odds ratios by comparing C/EBPβ ChIP-seq reads and input sample ChIP-seq reads. Y=0 indicates where TF binding has an expected amount of activity. P-values were based on two-sample t-tests by comparing the InDel groups of each test region. (**D**) Sequencing data of ER-HoxB8 cells at co-binding site of PU.1 and C/EBPβ. Highlighted is test region #6 whose DNA sequence from PU.1 motif to C/EBPβ motif is shown. (**E**) Log2 odds ratios of test regions #6 as a function of InDel size.

We tested six PU.1 and C/EBPβ co-binding sites with the original motif spacing ranging from 26 to 55 bp (Figure 5-table supplement 1) and quantified the effects of various InDels on C/EBPβ binding. Among the six test regions, three of them have supportive evidence from naturally occurring InDels of mouse strains (region #1, #3, #5) and the other three don’t (region #2, #4, #6). Based on the bioinformatic analysis of the ultra-deep sequencing reads from the input DNA samples, we saw that the CRISPR/Cas9 system generated a wide range of InDels with most deletions being < 30 bp and short insertions usually less than 5 bp (Fig. 5B). It provides longer deletions than natural genetic variations found across mouse strains (Figure 3-figure supplement 1) and in human populations (Figure 2A). After classifying ChIP-seq reads based on the InDel size and whether the InDel overlaps with any of the PU.1 and C/EBP motifs, we estimated the effect size of InDels on C/EBPβ binding by calculating the odds ratio between C/EBPβ ChIP-seq reads and input sample ChIP-seq reads for every InDel group. We found that InDels altering spacing have significantly weaker effects on C/EBPβ binding in comparison to those overlapping with at least one of the motifs (Fig. 5C). For some test regions, the effects of pure spacing alterations are almost negligible, exemplified by test region #6 (Fig. 5D, E) and test region #1 (Figure 5-figure supplement 1). Test region #6 is located near a highly expressed gene *Prdx1* and has strong binding of PU.1 and C/EBPβ binding and strong signals of H3K27ac and chromatin accessibility indicated by ATAC-seq in ER-Hoxb8 cells, which all support its potential regulatory function (Fig. 5D). The PU.1 and C/EBPβ motifs at this region are 26 bp apart. In general, spacing alterations ranging from 5-bp increase to 22-bp decrease did not have a strong effect on TF binding, indicated by a log2 odds ratio close to 0 (Fig. 5E). A small number of outliers were observed at each region where specific InDels resulted in substantial loss of binding (e.g., −20 bp, Fig. 5E). C/EBPβ binding at these specific InDels was generally discontinuous with 1-bp increments (e.g., −19 bp and −21 bp, Fig. 5E). The basis for these highly localized changes in the odds ratio in a small fraction of InDels that alter spacing is unclear. On the contrary, deletions overlapping with motifs resulted in a general decrease in TF binding activity. Similar results were found at test region #1 where PU.1 and C/EBPβ motifs are 41 bp apart (Figure 5-figure supplement 1A). This *Ly9* enhancer also has a 5-bp insertion between PU.1 and C/EBPβ motifs in BALB, NOD, and PWK mice, and shows unaffected binding of PU.1 and C/EBPβ in the BMDMs of these strains (Figure 5-figure supplement 1B). As a result of the synthetic InDels, the C/EBPβ binding activity was generally unaffected by spacing alterations only whereas deletions affecting motifs substantially diminished TF binding (Figure 5-figure supplement 1C).

## Discussion

By classifying the genome-wide spacing relationships of 75 co-binding TFs as “constrained” or “relaxed”, we revealed that relaxed spacing relationships were the dominant pattern of interaction for majority of these factors. Among these factors, approximately half could also participate in constrained spacing relationships with specific TF partners. We confirmed TF pairs known to exhibit constrained relationships (e.g., GATA1-TAL1) and identified previously unreported constrained relationships for additional pairs, including EGR1 and JUND. Overall, this finding of a subset of constrained TF interactions on a genome wide level is consistent with the locus-specific examples provided by functional and structural studies of the interferon-β enhanceosome (Panne, 2008) and in vivo studies of synthetically modified enhancer elements in Ciona (Farley et al., 2015). Each of these examples represents genomic regulatory elements in which key TF motifs are tightly spaced in their native contexts (i.e., 0-9 bp between motifs). Direct protein-protein interactions are observed between bound TFs at the interferon-β enhanceosome, analogous to interactions defined for cooperative TFs that form ternary complexes (Morgunova & Taipale, 2017; Reményi et al., 2003). In the present studies, InDels between TF pairs exhibiting constrained spacings were under large selective constraints that were comparable to InDels at motifs, suggesting a deleterious effect of these spacing alterations on TF binding. However, the spacing analyses in this study did not directly consider the possible overlap or lack of spacing between TF binding sequences. Thus, we are not able to clearly distinguish effects of spacing alterations from effects of InDels on motifs at sites of tightly spaced composite motifs.

The observation that most TF pairs exhibited relaxed spacing relationships has intriguing implications for the mechanisms by which functional enhancers and promoters are selected from chromatinized DNA. In contrast to ternary complexes of TFs that cooperatively bind to composite elements as a unit, relaxed spacing relationships appear to not require specific protein-protein interactions between TFs for collaborative binding at most genomic locations. Although pioneering TFs necessary for selection of cell-specific enhancers have been reported to recognize their motifs within the context of nucleosomal DNA (Zaret & Carroll, 2011), the basis for collaborative binding interactions between TFs with relaxed spacings remains poorly understood.

While the current studies relying on natural genetic variation and mutagenesis experiments concluded clear tolerance of spacing alterations between motifs of TFs with relaxed spacings, the extent to which this set of binding sites is representative of all regulatory elements is unclear. For example, we observed outliers in which significant differences in TF binding between mouse strains were associated with InDels occurring between motifs. However, the proportion of outliers was generally similar to that observed at genomic regions lacking such InDels, and such strain differences may be driven by distal effects of genetic variation on interacting enhancer or promoter regions (Hoeksema et al., 2021; Link, Duttke, et al., 2018). The remarkable tolerance of synthetic InDels at two independent endogenous genomic locations between PU.1 and C/EBPβ binding sites strongly support the generality of relaxed binding interactions for these two proteins. Intriguingly, while the densities of C/EBP motifs increase with decreasing distance to PU.1 motifs over a 100 bp range (Fig. 3A), deletions from 1 to >30 bp between PU.1-C/EBPβ pairs did not result in improved binding. Instead, relatively constant binding was observed with progressive deletions bringing two motifs close together until the deletions started to cause mutations in one or both motifs. This is consistent with the lack of correlation between DNA binding strengths and distances between these factors (Figure 3B). A limitation of these studies is that few and relatively short insertions were obtained, preventing conclusions as to the extent to which increases in spacing are tolerated.

In concert, the present studies provide a basis for estimation of the potential phenotypic consequences of naturally occurring InDels in non-coding regions of the genome. The majority of naturally occurring InDels are less than 5 bp in length. In nearly all cases, InDels of this size range between motifs for TFs that have relaxed binding relationships are unlikely to alter TF binding and function, and InDels of much greater length are frequently tolerated. In contrast, InDels between motifs for TFs that have constrained binding relationships have the potential to result in biological consequences. Application of these findings to the interpretation of non-coding InDels that are associated with disease risk will require knowledge of the relevant cell type in which the InDel exerts its phenotypic effect and the types of TF interactions driving the selection and function of the affected regulatory elements.

## Methods

### Sequencing data processing

We downloaded two replicates for each TF ChIP-seq data from ENCODE data portal (Davis et al., 2018). The mouse BMDM data and the human endothelial cell data were downloaded from the GEO database with accession number GSE109965 (Link, Duttke, et al., 2018) and GSE139377 (Stolze et al., 2020), respectively. We mapped ChIP-seq and ATAC-seq reads using Bowtie2 v2.3.5.1 with default parameters (Langmead & Salzberg, 2012) and mapped RNA-seq reads using STAR v2.5.3 (Dobin et al., 2013). All the human data downloaded from ENCODE were mapped to the hg38 genome. Data from C57BL/6J mice were mapped to the mm10 genome. Data from other mouse strains and endothelial cell data from different individuals were mapped to their respective genomes built by MMARGE v1.0 (Link, Romanoski, et al., 2018). More details are described below.

Based on the mapped ChIP-seq data, we called TF binding sites or peaks using HOMER v4.9.1 (Heinz et al., 2010). For data with replicates including ENCODE data and mouse data, we first called unfiltered 200-bp peaks using HOMER “findPeaks” function using parameters “-style factor −L 0 −C 0 -fdr 0.9 -size 200” and then ran IDR v2.0.3 with default parameters (Li et al., 2011) to obtain reproducible peaks. For data without replicates including human endothelial cell data and ER-HoxB8 ChIP-seq data, we called peaks using HOMER “findPeaks” with the default setting and parameters “-style factor -size 200”.

Activity of TF binding was quantified by the ChIP-seq tag counts within 300-bp around peak centers and normalized by library size using HOMER “annotatePeaks.pl” script with parameters “-norm 1e7 -size −150,150”. Activity of promoter and enhancer was quantified by normalized H3K27ac ChIP-seq tags within 1,000-bp regions around TF peak centers using parameters “- norm 1e7 -size −500,500”.

### Motif identification

Based on DNA sequences of the TF binding sites, we calculated motif scores by the dot products between PWMs and sequence vectors using Biopython package (Cock et al., 2009). The PWMs were obtained from either the JASPAR database (Fornes et al., 2020) or de novo motif analysis using HOMER “findMotifsGenome.pl” script (Heinz et al., 2010) if unavailable in the JASPAR database. The original PWMs were then trimmed to keep only the core motifs starting from the first position where information content greater than 0.3 to the last position of information content greater than 0.5 (Ng et al., 2014). The valid motifs were identified by a motif score passing a false positive rate (FPR) 0.1% and a location within 50 bp close to the peak center. The motif spacing is computed as the edge-to-edge distance between two core motifs at TF cobinding sites. If there are multiple valid motifs for one or both TFs, we computed the spacing between all possible combinations of valid motifs.

### Characterization of spacing relationships

To test for the constrained spacing relationship between any two TFs, we developed a method to identify “spikes” in the spacing distribution. We first counted the TF pair distances at single-base-pair resolution ranging from −100 bp to +100 bp. Next, we computed the slope at each position using the following formula:

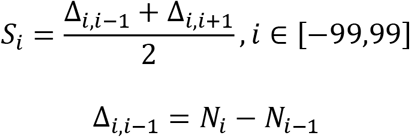

*S_i_* is the average of single-step forward and backward slope at position *i*. *N_i_* represents the number of TF pair at position *i*, and Δ is the difference in the number of TF pairs between two locations. We conducted permutation tests to compare each *Si* to a simulated null distribution to determine a p-value based on the percentile rank. P-value smaller than 6.25e-05 (familywise error rate=0.05/200/4) is called significant, indicating a spike is found among motif spacing between the testing TF pair. The null distribution was generated by 1,000 iterations of 1,000 random spacing between 0 and 100 bp.

To test for the relaxed spacing relationship, we used Kolmogorov–Smirnov (KS) test to compare a spacing distribution to the random distribution. We randomly sampled integers between −100 and 100 to match the same size of the testing spacing distribution and then tested the spacing distribution against the distribution of the random integers to obtain a p-value. We repeated the above process 100 times and reported the average p-value.

### Categorization of gnomAD variants based on allele frequency

We obtained InDels from gnomAD v3.1 (Karczewski et al., 2020). These gnomAD variants were overlapped with TF co-binding sites, specifically with two TF motifs and their intermediate sequences. For TF pairs with constrained spacing relationships, we only kept the co-bindings that have the significant constrained spacing +/− 2 bp. To account for region-by-region variation in selective pressure, we also overlapped variants with 100-bp upstream and 100-bp downstream background regions outside of TF co-binding sites. For each co-binding site, we categorized InDels into high-frequency variants (AF > 0.01%), rare variants (AF < 0.01%, AC > 1), and singletons (AC = 1) and computed the odds ratios of different categories of InDels between motif or intermediate regions and background regions.

### Genetic variation processing and genome building

Genetic variation of the five mouse strains was obtained from (Keane et al., 2011), and that of the human individuals from which endothelial cell data were generated was derived from (Stolze et al., 2020). We used MMARGE v1.0 with default variant filters (Link, Romanoski, et al., 2018) to build separate genomes for each mouse strain and human individual. The sequencing data from different samples were respectively mapped to the corresponding genomes and were then shifted to a common reference genome using MMARGE “shift” function to facilitate comparison at homologous regions. The reference genome is mm10 for mouse strains and hg19 for human individuals.

### Motif mutation analysis

We used MAGGIE (Shen et al., 2020) to identify functional motifs for different TF binding. To prepare the inputs into MAGGIE based on the mouse strains data, we adapted a similar strategy as described in (Shen et al., 2020). In brief, we conducted pairwise orthogonal comparisons of TF peaks between each possible pair of the five strains to find strain-differential peaks. We then extracted pairs of 200-bp sequences around the centers of the differential peaks from the genomes of two comparative strains, the ones with TF binding as positive sequences paired with those without TF binding as negative sequences. For the QTLs of human endothelial cells, MAGGIE can directly work with a VCF file of QTLs with effect size and effect direction indicated in a column of the file. We ran MAGGIE separately for each type of QTLs and reported the significant motifs together with their p-values, which passed false discovery rate (FDR) < 0.05 after the Benjamini–Hochberg controlling procedure.

### Categorization of genetic variation based on impacts on motif or spacing

We categorized genetic variation based on its impact on motif affinity and motif spacing. Motif mutations were defined by at least 2-bit difference in the motif score, which is equivalent to approximately 4-fold difference in the binding likelihood. Mutations of other functional motifs identified by MAGGIE required that at least one of the functional motifs had motif mutations. InDels were first classified into motif mutation categories if eligible before being considered in motif spacing. Therefore, spacing alterations were InDels between target motifs without any motif mutations. Variants fitting neither motif mutation nor spacing alteration were gathered in a separate group as a control. Another control category during analysis of mouse strains data was defined by TF binding sites that have no genetic variation between strains.

### Statistical testing of effect size

We conducted Mann-Whitney U tests between the control category and one of the testing categories to test for significance. We also obtained the Cohen’s *d* (Sullivan & Feinn, 2012) between the control category and the testing categories and Spearman’s correlation coefficients as measures of effect size.

### ER-HoxB8 cell-derived macrophage culture and CRISPR knockout

Bone marrow cells were isolated from femurs and tibias of a Cas9-expressing transgenic mouse (Jackson Laboratory, No.028555). Murine stem cell virus-based expression vector for ER-HoxB8 was gifted from Dr. David Sykes (Massachusetts General Hospital, Boston, MA). Cas9-expressing ER-HoxB8 conditionally immortalized myoid progenitor cells were generated following established protocols (Wang et al., 2006). In brief, bone marrow cells were purified with a Ficoll gradient (Ficoll-Paque-Plus, Sigma-Aldrich) and resuspended in RPMI 1640 containing 10% FBS, 1% penicillin/streptomycin and 10 ng/ml each of SCF, IL-3 and IL-6 (PeproTech). After 48 hours culture, 2.5×10^5^cells in 1 ml were transduced with 2 ml of ER-HoxB8 retrovirus (in DMEM with 30% FBS) containing 0.5 μl/ml lentiblast A (OZ Biosciences), 2.5 μl/ml lentiblast B (OZ Biosciences) and 8 μg/ml polybrene (Sigma-Aldrich) in a well of fibronectin (Sigma-Aldrich)-coated 6-well culture plates and centrifuged at 1000g for 90 min at 22°C. After transduction, 6 ml of ER-HoxB8 cell culture media (RPMI 1640 supplemented with 10% FBS, 1% penicillin/streptomycin, 0.5 μM β-estradiol (Sigma-Aldrich), and 20 ng/ml GM-CSF (PeproTech)) were added and an additional half-media exchange with ER-Hoxb8 media performed the next day. Transduced cells were selected with G418 (Thermo Fisher) at 1 mg/ml for 48 hours. Thereafter, cells were maintained in ER-HoxB8 cell culture media. For baseline ATAC-seq and ChIP-seq of ER-HoxB8 cells prior to gRNA transduction, cells were washed twice with PBS, plated at a density of 3×10^6^cells per 10 cm culture plate, and differentiated into macrophages in DMEM with 10% FBS, 1% pencillin/streptomycin, and 17 ng/ml M-CSF (Shenandoah) for 7 days with 2 culture media exchanges. Differentiated cells were washed twice with PBS and collected for sequencing experiments.

Guide RNA lentiviruses were prepared as previously described (Fonseca et al., 2019) with modifications as follows. LentiGuide-mCherry was generated by modifying lentiGuide-puro (Addgene) to remove a puromycin-resistant gene and replace it with mCherry. gRNA sequences directed between the PU.1 and C/EBP motifs (and one each directed towards the motif itself) were designed with CHOPCHOP web tool for genome engineering (Labun et al., 2019). One CRISPR gRNA oligonucleotide was inserted for each target via PCR into a BsmBI cleavage site. A list of gRNA targets used in this article is shown in Figure 5-table supplement 1. Lenti-X 293T cells (Clontech) were seeded in poly-D-lysin (Sigma-Aldrich) coated 10 cm tissue culture plates at a density of 3.5 million cells per plate in 10 ml of DMEM containing 10% FBS and 1% penicillin/streptomycin, and then incubated overnight at 37°C. After replacement of the media to 6 ml of DMEM containing 30% FBS, plasmid DNAs (5 μg of lentiviral vector, 3.75 μg of psPAX2 and 1.25 μg of pVSVG) were transfected into LentiX-293T cells using 20 μl of X-tremeGENE™ HP DNA Transfection Reagent (Roche) at 37°C overnight. The media was replaced with DMEM containing 30% FBS and 1% penicillin/streptomycin, and then cultured at 37°C overnight. The supernatant was filtrated with 0.45 μm syringe filters and used as lentivirus media. Cell culture media was replaced, and virus was collected again after 24 hours. 1×10^6^ Cas9-expressing ER-HoxB8 cells were transduced with virus in 2 ml of lentivirus media and 1 ml of ER-HoxB8 cell media containing 0.5 μl/ml lentiblast A, 2.5 μl/ml lentiblast B, and 8 μg/ml polybrene in a well of fibronectin-coated 6-well culture plates and centrifuged at 1000 g for 90 min at 22°C. After the transduction, 6 ml of ER-HoxB8 cell media was added to each well. Half of the media was exchanged the next day and in the following days, cells were expanded and passaged into T75 flasks. After 5 days, 250,000 successfully transduced cells (indicated by mCherry fluorescence) for each gRNA were sorted by FACS using a Sony MA900. After FACS, cells were expanded in ER-HoxB8 culture media. Differentiation into macrophages was carried out as above in DMEM supplemented with M-CSF.

### RNA-seq library preparation

Total RNA was isolated from cells and purified using Direct-zol RNA Microprep columns according to the manufacturer’s instructions (Zymo Research). 500 ng of total RNA were used to prepare sequencing libraries from polyA enriched mRNA as previously described (Link, Duttke, et al., 2018). Libraries were PCR-amplified for 14 cycles, size selected using Sera-Mag Speedbeads (Thermo Fisher Scientific), quantified by Qubit dsDNA HS Assay Kit (Thermo Fisher Scientific) and 75-bp single-end sequenced on a HiSeq 4000 (Illumina).

### ATAC-seq library preparation

ATAC-seq libraries were prepared as previously described (Hoeksema et al., 2021). In brief, 5×10^5^ cells were lysed at room temperature in 50 μl ATAC lysis buffer (10 mM Tris-HCl, pH 7.4, 10 mM NaCl, 3 mM MgCl2, 0.1% IGEPAL CA-630) and 2.5 μL DNA Tagmentation Enzyme mix (Nextera DNA Library Preparation Kit, Illumina) was added. The mixture was incubated at 37°C for 30 minutes and subsequently purified using the ChIP DNA purification kit (Zymo Research) as described by the manufacturer. DNA was amplified using the Nextera Primer Ad1 and a unique Ad2.n barcoding primers using NEBNext High-Fidelity 2X PCR MM for 8-14 cycles. PCR reactions were size selected using TBE gels for 175 – 350 bp and DNA eluted using gel diffusion buffer (500 mM ammonium acetate, pH 8.0, 0.1% SDS, 1 mM EDTA, 10 mM magnesium acetate) and purified using ChIP DNA Clean & Concentrator (Zymo Research). Samples were quantified by Qubit dsDNA HS Assay Kit (Thermo Fisher Scientific) and 75-bp single-end sequenced on HiSeq 4000 (Illumina).

### Crosslinking for ChIP-seq

For PU.1, C/EBPβ, and H3K27ac ChIP-seq, culture media was removed, and plates were washed once with PBS and then fixed for 10 minutes with 1% formaldehyde (Thermo Fisher Scientific) in PBS at room temperature. Reaction was then quenched by adding glycine (Thermo Fisher Scientific) to 0.125M. After fixation, cells were washed once with cold PBS and then scraped into supernatant using a rubber policeman, pelleted for 5 minutes at 400x*g* at 4°C. Cells were transferred to Eppendorf DNA LoBind tubes and pelleted at 700x*g* for 5 minutes at 4°C, snap-frozen in liquid nitrogen and stored at −80°C until ready for ChIP-seq protocol preparation.

### Chromatin immunoprecipitation

Chromatin immunoprecipitation (ChIP) was performed in biological replicates as described previously (Hoeksema et al., 2021). Samples were sonicated using a probe sonicator in 500 μl lysis buffer (10 mM Tris/HCl pH 7.5, 100 mM NaCl, 1 mM EDTA, 0.5 mM EGTA, 0.1% deoxycholate, 0.5% sarkozyl, 1x protease inhibitor cocktail). After sonication, 10% Triton X-100 was added to 1% final concentration and lysates were spun at full speed for 10 minutes. 1% was taken as input DNA, and immunoprecipitation was carried out overnight with 20 μl Protein A Dynabeads (Invitrogen) and 2 μg specific antibodies for C/EBPβ (Santa Cruz, sc-150), PU.1 (Santa Cruz, sc-352X), and H3K27ac (Active Motif, 39135). Beads were washed three times each with wash buffer I (20 mM Tris/HCl, 150 mM NaCl, 0.1% SDS, 1% Triton X-100, 2 mM EDTA), wash buffer II (10 mM Tris/HCl, 250 mM LiCl, 1% IGEPAL CA-630, 0.7% Na-deoxycholate, 1 mM EDTA), TE 0.2% Triton X-100 and TE 50 mM NaCl and subsequently resuspended 25 μl 10 mM Tris/HCl pH 8.0 and 0.05% Tween-20. ChIP-seq libraries were prepared on the Dynabeads as described below. For locus specific enrichment ChIP-seq, bead complex was resuspended in 50 μl 1% SDS-TE. 4 μl ProtK, 4 μl RNase A, 3 μl 5 M NaCl was added to these and the input samples and incubated at 50°C for 1 hour, reverse crosslinked at 65°C overnight and then eluted from the beads.

### ChIP-seq library preparation

ChIP libraries were prepared while bound to Dynabeads using NEBNext Ultra II Library preparation kit (NEB) using half reactions. DNA was polished, polyA-tailed and ligated after which dual UDI (IDT) or single (Bioo Scientific) barcodes were ligated to it. Libraries were eluted and crosslinks reversed by adding to the 46.5 μl NEB reaction 16 μl water, 4 μl 10% SDS, 4.5 μl 5 M NaCl, 3 μl 0.5 M EDTA, 4 μl 0.2 M EGTA, 1 μl RNAse (10 mg/ml) and 1 μl 20 mg/ml proteinase K, followed by incubation at 55°C for 1 hour and 75°C for 30 minutes in a thermal cycler. Dynabeads were removed from the library using a magnet and libraries were cleaned up by adding 2 μl SpeedBeads 3 EDAC (Thermo) in 124 μl 20% PEG 8000/1.5 M NaCl, mixing well, then incubating at room temperature for 10 minutes. SpeedBeads were collected on a magnet and washed two times with 150 μl 80% ethanol for 30 seconds. Beads were collected and ethanol removed following each wash. After the second ethanol wash, beads were air dried and DNA eluted in 12.25 μl 10 mM Tris/HCl pH 8.0 and 0.05% Tween-20. DNA was amplified by PCR for 14 cycles in a 25 μl reaction volume using NEBNext Ultra II PCR master mix and 0.5 μM each Solexa 1GA and Solexa 1GB primers. Libraries were size selected using TBE gels for 200 – 500 bp and DNA eluted using gel diffusion buffer (500 mM ammonium acetate, pH 8.0, 0.1% SDS, 1 mM EDTA, 10 mM magnesium acetate) and purified using ChIP DNA Clean & Concentrator (Zymo Research). Sample concentrations were quantified by Qubit dsDNA HS Assay Kit (Thermo Fisher Scientific) and 75-bp single-end sequenced on HiSeq 4000.

### Biotin-mediated locus specific enrichment ChIP-seq library preparation

After performing the target-specific ChIPs, we performed an initial PCR for locus-specific amplicon enrichment using NEBNext 2X High Fidelity PCR MM (NEB) and 5’-biotinylated stub adapter primers specific to appropriate genomic regions to be interrogated (Figure 5-table supplement 1). Initial hotstart/denaturation at 98°C for 30 sec was followed by 10 cycles of amplification (98°C for 15 sec, 65-67°C for 15 sec, 72°C for 30 sec) and then a final elongation at 72°C for 5 min. After this, we performed a 0.7X AmpureXP clean-up and eluted in 20 μl 0.5x TT (5 mM Tris pH 8.0 + 0.025% Tween20). Dynabeads MyOne Streptavidin T1 beads were then washed in 1x Wash Binding Buffer (WBB, 2X WBB: 10 mM Tris-HCl (pH 7.5), 1 mM EDTA, 2 M NaCl, 0.1% Tween) and resuspended beads at 20 μl per sample in 2x WBB. 20 μl prepared Dynabeads MyOne Streptavidin T1 beads (in 2x WBB) were then added to cleaned up 20 μl 0.5x TT PCR fragments, mixed and incubated for 60min at RT with mild shaking. After this, beads were collected on a magnet and washed twice with 150 μl 1x WBB and once with 180 μl TET (TE + 0.05% Tween-20). Finally, beads were resuspended in 25 μl 0.5x TT and on bead PCR for addition of Illumina-specific adapters and 10-bp Unique Dual Indexes (UDIs) using NEBNext 2X High Fidelity PCR MM (NEB) and 25 PCR cycles was performed (Figure 5-table supplement 2). Libraries were size selected using TBE gels for 300-500 bp and DNA eluted using gel diffusion buffer and purified using ChIP DNA Clean & Concentrator (Zymo Research). Samples were quantified by Qubit dsDNA HS Assay Kit (Thermo Fisher Scientific) and 150-bp paired-end sequenced on NextSeq 500 (Illumina).

### Analysis of variable InDels from CRISPR experiments

We mapped the reads to the target regions using the local alignment mode of Bowtie2 v2.3.5.1 (Langmead & Salzberg, 2012). To allow for InDels with tens of bases, we reduced the gap extend penalty and increased the gap open penalty so that the gaps could be long but not occur at multiple locations. Here are the adjusted parameters used in our mapping process: --local --rdg 10,1 --rfg 10,1. The mapped reads with gaps or InDels at unexpected locations rather than the Cas9 cut sites were removed. This step filtered out approximately 1% of the total reads (Figure 5-table supplement 2). The remaining reads were grouped based on the InDel size and whether the InDel overlaps with any of the PU.1 and C/EBP motifs. Tag counts were used as quantification of the signal intensity. InDel groups with less than 0.05% of the input reads were filtered out to reduce the low-intensity data. The effect of each InDel group on TF binding was computed by the odds ratio between TF ChIP-seq tags and input sample ChIP-seq tags: (# TF tags for an InDel group / # the rest of TF tags) / (# input tags for the same InDel group / # the rest of input tags).

## Data and code availability

All sequencing data generated from this study have been made available by deposition in the GEO database: GSE178080. All raw tag counts are available in Figure 5-source data 1. The UCSC genome browser was used to visualize sequencing data. The codes for data analysis and the processed files of ENCODE data are available on our Github repository: https://github.com/zeyang-shen/spacing_pipeline.

## Acknowledgements

The authors would like to thank J. Collier for technical assistance, the IGM core for library sequencing, L. Van Ael for assistance with manuscript preparation. These studies were supported by NIH grants DK091183 and HL147835 and a Leducq Transatlantic Network grant 16CVD01 to CKG. TAP was supported by NIH grant T32DK007044. MAH was supported by a Rubicon grant from the Netherlands Organization for Scientific Research and postdoctoral grants from the Amsterdam Cardiovascular Sciences Institute and the American Heart Association.

