## Supplemental figures for "Systematic analysis of naturally occurring insertions and deletions that alter transcription factor spacing identifies tolerant and sensitive transcription factor pairs"

### Supplementary figures

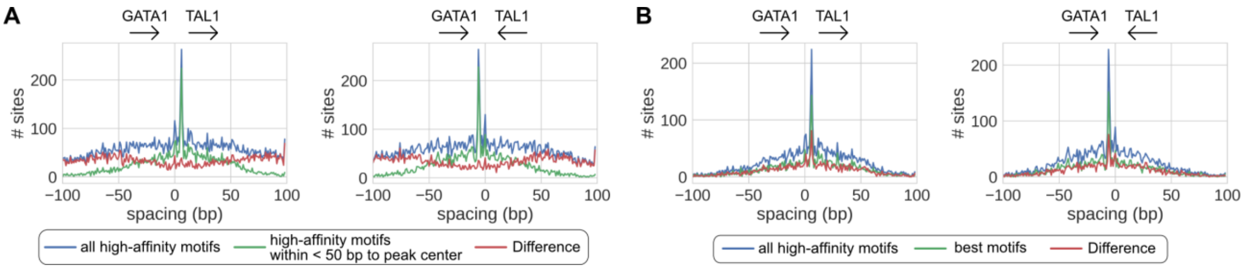


**Figure 1 – figure supplement 1**. Effects of different motif scanning criteria. (A) Motifs proximal to peak centers are potentially more confident than motifs distal from peak centers. (B) All motifs passing FPR < 0.001 are potentially as confident as the best motif of every peak.

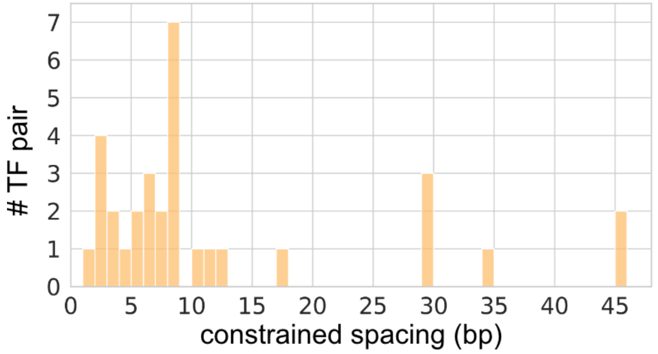


**Figure 1 - figure supplement 2**. Constrained spacings for the significant TF pairs with constrained spacing relationships.

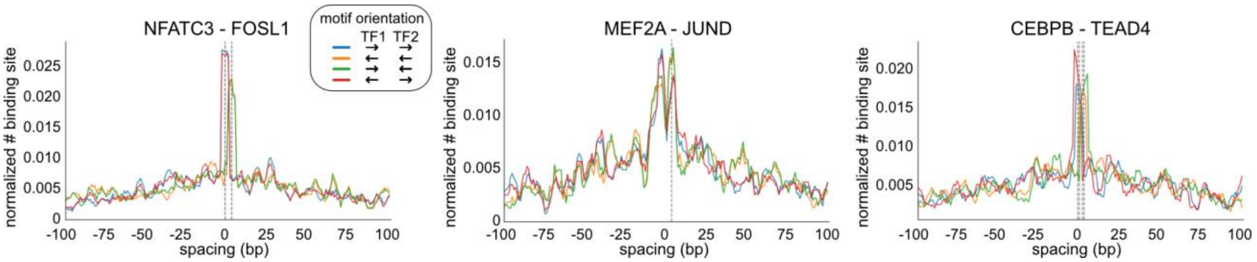


**Figure 1 - figure supplement 3**. Examples of TF pairs with constrained spacing relationships.

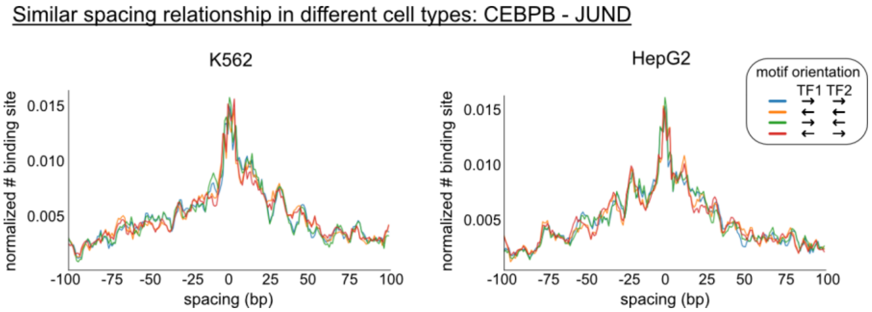


**Figure 1 - figure supplement 4**. Comparison of the spacing relationships of same TF pairs in different cell types.

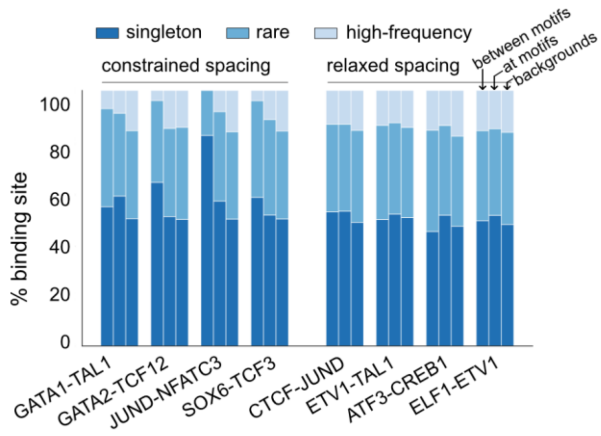


**Figure 2 - figure supplement 1**. Composition of InDels with different allele frequency for representative TF pairs. InDels were divided into high-frequency variants (AF > 0.01%), rare variants (AF < 0.01%, AC > 1), and singletons (AC = 1).

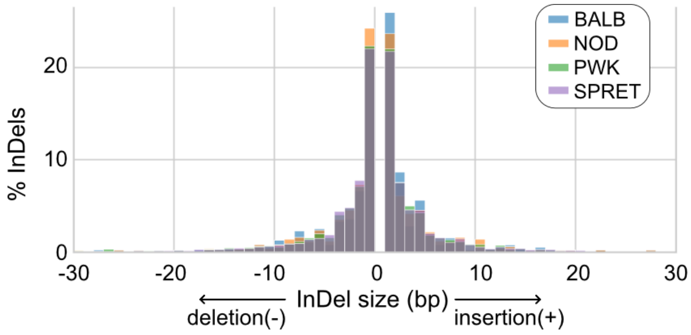


**Figure 3 - figure supplement 1**. Size distributions of InDels at PU.1 and C/EBPβ co-binding sites across mouse strains.

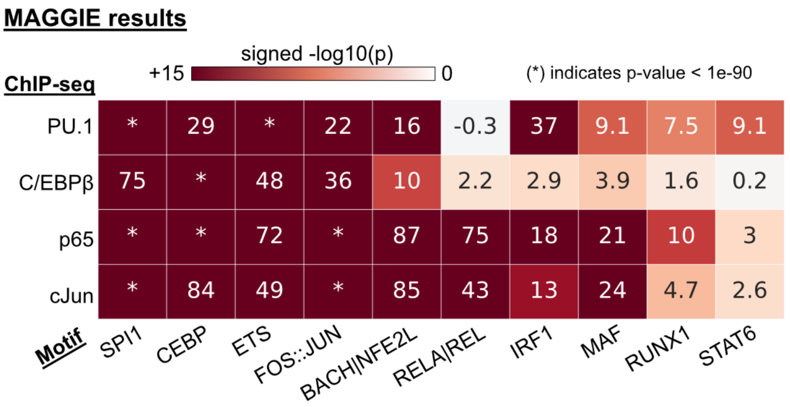


**Figure 3 - figure supplement 2**. Functional motifs identified by MAGGIE for different TF binding.

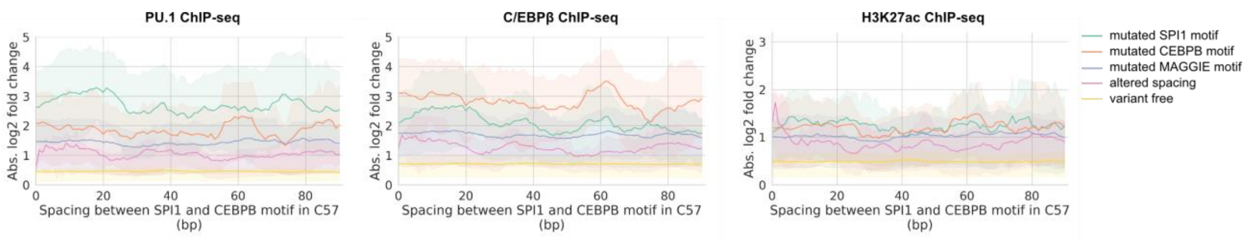


**Figure 3 - figure supplement 3.** Absolute log2 fold changes of ChIP-seq tags in relationship with the initial spacing between PU.1 and C/EBPβ motif in the reference mm10 genome. The results were aggregated from all four pairwise comparisons.

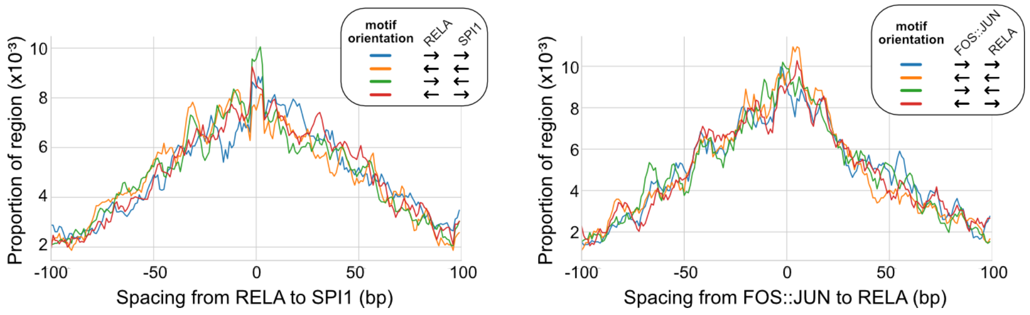


**Figure 3 - figure supplement 4**. Spacing distributions between LDTFs and SDTFs. Left: p65 and PU.1. Right: p65 and cJun.

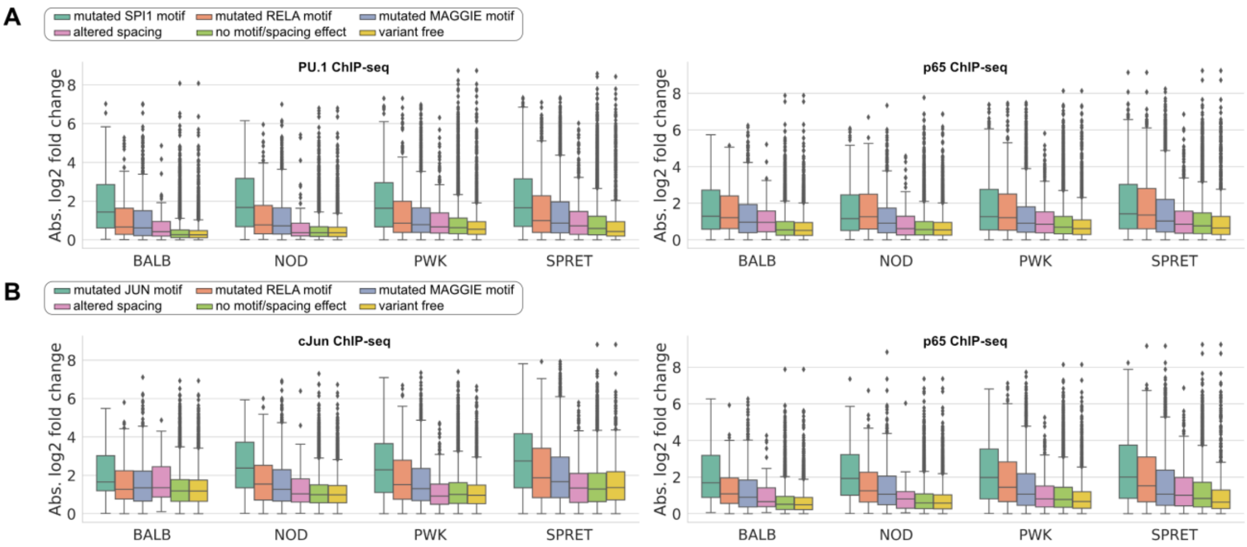


**Figure 3 - figure supplement 5**. Absolute log2 fold changes of ChIP-seq tags between C57 and another strain for LDTFs and SDTFs. (A) PU.1 and p65 binding at their co-binding sites and (B) cJun and p65 binding at their co-binding sites.

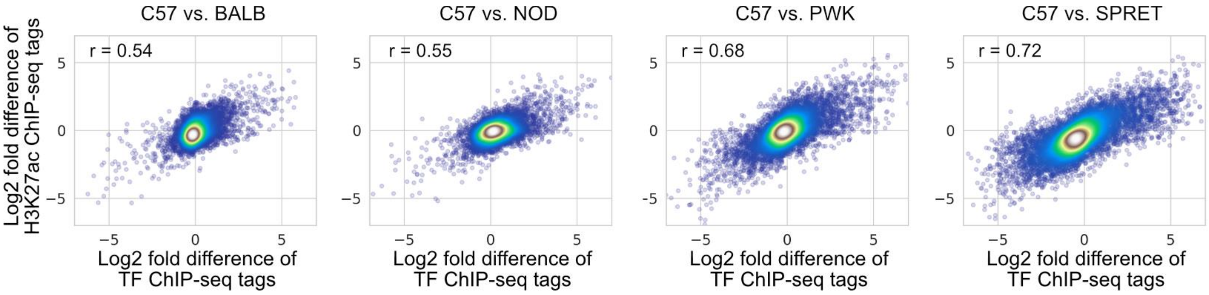


**Figure 3 - figure supplement 6**. Correlations between changes in TF binding activity and changes in the H3K27ac level.

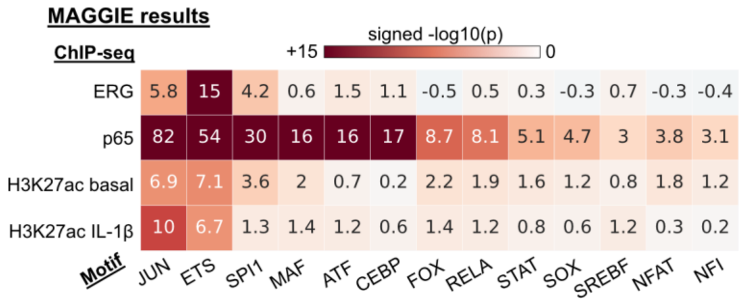


**Figure 4 - figure supplement 1**. Functional motifs identified by MAGGIE based on bQTLs.

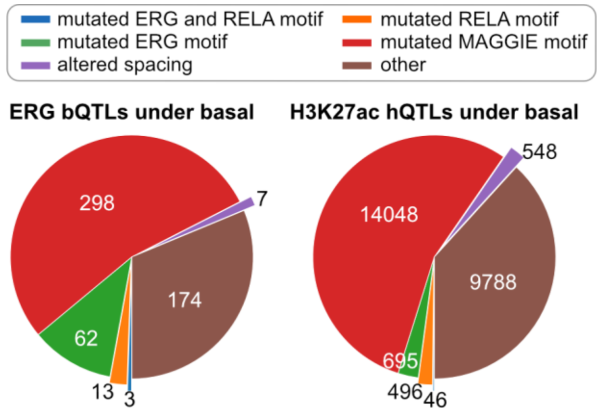


**Figure 4 - figure supplement 2**. Classification of chromatin QTLs based on the effects on motif and spacing for basal condition.

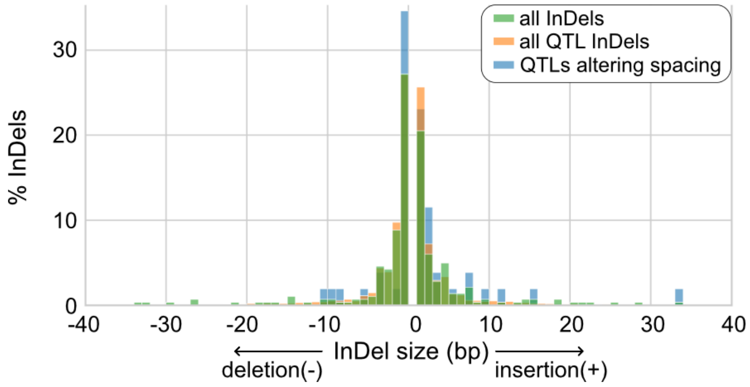


**Figure 4 - figure supplement 3**. Size distributions of InDels from human endothelial cell donors.

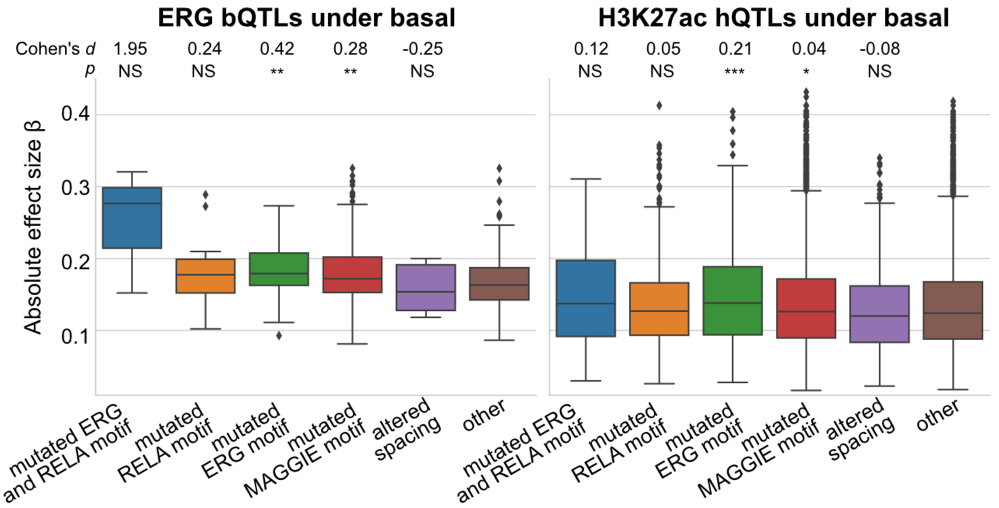


**Figure 4 - figure supplement 4**. Absolute correlation coefficients of different QTLs for basal condition. Cohen's *d* and Mann-Whitney U test p-values comparing against the "other" group are displayed on top. $^{*}p<0.01$, $^{**}p<0.001$, $^{***}p<0.0001$.

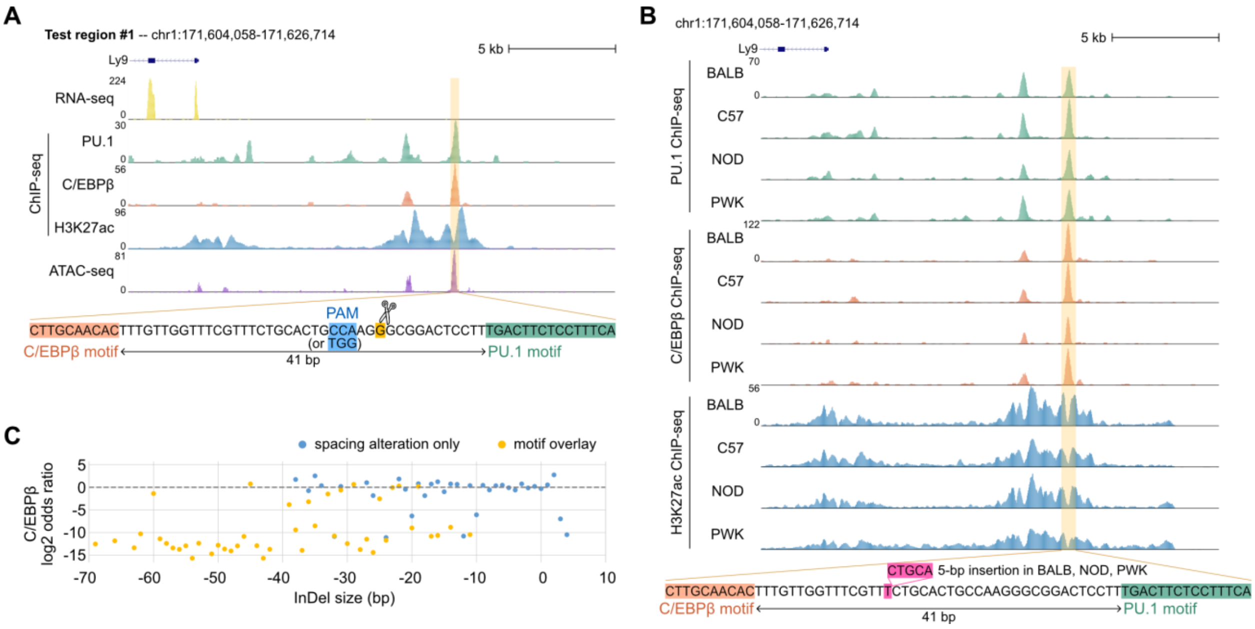


**Figure 5 - figure supplement 1**. (A) Mouse strains data for test region #1. (B) Sequencing data of ER-HoxB8 cells for test region #1. (C) Log2 odds ratios of test regions #1 as a function of InDel size.
